# Adipose deficiency and aberrant autophagy in a *Drosophila* model of MPS VII is corrected by pharmacological stimulators of mTOR

**DOI:** 10.1101/2021.09.05.459029

**Authors:** Indrani Basu, Sudipta Bar, Mohit Prasad, Rupak Datta

## Abstract

Mucopolysaccharidosis type VII (MPS VII) is a recessively inherited lysosomal storage disorder caused due to β-glucuronidase (β-GUS) enzyme deficiency. Prominent clinical symptoms include hydrops fetalis, musculoskeletal deformities, neurodegeneration and hepatosplenomegaly leading to premature death in most cases. Apart from these, MPS VII is also characterized as adipose storage deficiency disorder although the underlying mechanism of this lean phenotype in the patients or β-GUS-deficient mice still remains a mystery. We addressed this issue using our recently developed *Drosophila* model of MPS VII (the CG2135^-/-^ fly), which also exhibited a significant loss of adiposity. We report here that the lean phenotype of the CG2135^-/-^ fly is due to fewer number of adipocytes, smaller lipid droplets and reduced adipogenesis. Our data further revealed that there is an abnormal accumulation of autophagosomes in the CG2135^-/-^ larvae due to autophagosome-lysosome fusion defect. Decreased lysosome-mediated turnover also led to attenuated mTOR activity in the CG2135^-/-^ flies. Interestingly, treatment of the CG2135^-/-^ larvae with mTOR stimulators, 3BDO or glucose, led to the restoration of mTOR activity with simultaneous correction of the autophagy defect and adipose storage deficiency. Our finding thus established a hitherto unknown mechanistic link between autophagy dysfunction, mTOR downregulation and reduced adiposity in MPS VII.

## 1. Introduction

Mucopolysaccharidosis type VII (MPS VII) or Sly syndrome is an autosomal recessive disorder caused due to deficiency of the lysosomal enzyme β-glucuronidase (β-GUS). This results in abnormal accumulation of undegraded and partially degraded glycosaminoglycans (dermatan sulfate, keratan sulfate, chondroitin sulfate) within the lysosomes[1]. Primary symptoms of MPS VII include hydrops fetalis, musculoskeletal deformities, hepatosplenomegaly, corneal clouding, mental retardation and heart, pulmonary and renal dysfunction subsequently leading to premature death [1–3]. MPS VII patients, females and males alike, were reported to be underweight with low body mass index (BMI) although the underlying cause of this lean feature remains unknown [3]. Interestingly, the bodyweight of the β-GUS deficient MPS VII mouse was also significantly less than normal and they were devoid of any visible white adipose tissue, a typical characteristic of adipose storage deficiency disorder [4]. It is important to note that such lean phenotype and decreased adiposity were found to be a common phenotype in mouse models of other many lysosomal storage disorders (LSDs) such as MPSI, MPSIIIB, Niemann-Pick type A/B and infantile neuronal ceroid lipofuscinosis [5].

Adipocytes are the body’s major reservoirs of energy that are stored predominantly in the form of lipid droplets obtained either directly from the diet or synthesized *de novo*. Adipogenesis is regulated by various transcription factors. Aberrant expression of these genes affects fat cell development and often leads to altered lipid homeostasis [6–8]. In 2007, Wolosynek *et al*. systematically studied the cause of adipose deficiency in several murine models of LSDs and concluded that diminished adiposity in these mice is not due to reduced food intake, lipid malabsorption, or higher caloric utilization [5]. Instead, they proposed that impaired recycling of the lysosomal precursors in these mice is due to autophagy defects that might lead to the breakdown of stored fat for energy generation required for *de novo* synthesis of raw materials. In a follow-up study, they tested this hypothesis by feeding the MPS VII mice with a nutrient-rich high calory diet in an attempt to reverse their lean phenotype. However, contrary to their expectation, the high-fat dietary intervention failed to completely restore the weight loss in these mice [9]. Thus, it may be concluded that altered energy balance owing to autophagy defect is not the primary reason for adipose deficiency observed in the MPS VII mice.

Functional crosstalk between the autophagy pathway and the adipogenic differentiation program was proposed by independent groups of researchers [10–13]. Adipose tissue-specific knockout of the atg7 gene exhibited a lean phenotype in mice with a stark reduction (∼20%) in the total white adipose tissue mass as compared to their wildtype counterparts. The white adipogenic program was shown to be repressed in these mice which promoted browning and beige phenotype of adipose tissues [13]. Atg5 and Atg7 knockdown in 3T3-L1 fibroblast cells also inhibited lipid accumulation and decreased expression of proteins responsible for adipogenesis [12]. All these reports pointed towards an important role of the autophagy process in adipocyte development.

Taking lead from these studies, we attempted to elucidate the underlying mechanism of reduced adiposity in MPS VII. We were intrigued by the question of whether autophagy defect could result in reduced adipogenesis in MPS VII. To address this, we performed extensive experiments using our recently developed *Drosophila* model of MPS VII where the CG2135 gene (the fly β-GUS homolog) was knocked out by genetic recombination [14]. Our studies revealed that MPS VII fly indeed suffers from adipose deficiency due to fewer numbers of adipocytes and reduced adipogenesis. These flies also exhibited an increased abundance of autophagosomes because of autophagosome-lysosome fusion defect in the larval fat body suggesting a causal relationship between reduced adipogenesis and aberrant autophagy. This notion was further strengthened when we found that the activity of the autophagy regulator mTOR was significantly downregulated in the MPS VII flies and we could correct both autophagy defect as well as adipose deficiency in these flies upon feeding them with mTOR stimulators (glucose and 3BDO). Taken together, this study provided important mechanistic insights into the adipose deficiency phenotype in MPS VII disease. This in turn allowed us to conceptualize corrective pharmacological intervention to restore normal adiposity in the MPS VII flies.

## 2. Materials and Methods

Unless otherwise specified, all reagents were purchased from Sigma-Aldrich (St. Louis MO). All oligonucleotide primers were procured from Integrated DNA Technologies and the sequence details are provided in Table S1.

### 2.1. *Drosophila* strains and maintenance

All *Drosophila* stocks were obtained from Bloomington *Drosophila* Stock Centre at Indiana University, USA. The following stocks were used: W^1118^, W;Actin GAL4/Cyo, W^1118^; Pin/Cyo; TM6Be/TM2ubx and y[1]w[1118];P{w[+mC]=UASp-GFP-mCherry-Atg8a}2 [15]. The CG2135^-/-^ fly construct used, was generated as described in Bar et al., 2018 [14]. The flies were maintained at a temperature of 25°C under constant humidity and controlled density with a 12-hour day-night cycle on a standard *Drosophila* food (cornmeal media containing dry yeast:15g/dL, cornflour:75g/dL, sugar:80g/dL). For larvae collection around 300 adult flies, in 3:2 female-male ratios were kept in embryo collection cage with 90mm dish containing the fly food. Wandering 3^rd^ instar larvae were obtained within 120 hrs A.E.L (after egg-laying). The larvae were collected using a fine paintbrush and the respective assays were performed both for WT and CG2135^-/-^ flies.

### 2.2. RNA isolation and qRT-PCR

Total RNA was isolated from 30 adult flies (male and female fly mixture) both of WT (W^1118^) and CG2135^-/-^ using TRIzol reagent following manufacturer’s instruction (Thermo Fischer Scientific). DNAse I (Invitrogen) treatment was done to remove DNA contaminants. 1µg of DNA-free RNA was reverse transcribed using a cDNA synthesis kit (Thermo Fischer Scientific). Transcript levels of different genes were quantified using this cDNA as the template for a specific set of primers as tabulated in Table S1. Real time PCR (qRT-PCR) was performed with SYBR Green PCR reaction mix (BioRad) using ABI Fast 7500 real-time PCR system (Applied Biosystems) in 20μl reaction volume. Normalization of target gene expression was done using rp49 as the reference gene (endogenous control). The results of quantitative real-time PCR were analyzed by the comparative threshold cycle (CT) method [16].

### 2.3. Isolation of Genomic DNA from *Drosophila*

For genomic DNA isolation from *Drosophila*, 15-20 flies were anaesthetized on ice followed by homogenization in 400µl of Buffer A (100mM Tris-Cl (pH=7.5), 100mM EDTA, 100mM NaCl and 0.5% SDS). The samples were incubated for 30mins at 65°C. 800μl of Buffer B (200ml potassium acetate (5M) and 500ml Lithium chloride 6M) was then added to each sample, mixed well multiple times followed by incubation on ice for 45mins to 1hr. The samples were then centrifuged at 12000rpm for 15mins at room temperature. An equal volume of phenol: chloroform was added and centrifuged at 12000 rcf for 15 mins at 4°C. 1ml of the supernatant was added to a fresh microcentrifuge tube and then mixed with 600μl of isopropanol. The following mixture was spun at 12000rpm for 15mins at room temperature. Finally, the supernatant was discarded and the pellet was dissolved in 70% ethanol (Merck), followed by air-drying and then resuspending in autoclaved water. The eluted DNA was used as the template for validation of transgenic construct flies by genomic DNA-PCR using primer sets as tabulated in Table S1.

### 2.4. Western blot

*Drosophila* larval lysate was prepared by homogenization of 30 larvae (each of starved WT, CG2135^-/-^, glucose and 3-BDO fed CG2135^-/-^) with a sterile Dounce homogenizer in 150µl of lysis buffer (25 mM Tris-Cl, pH-7.2, 140 mM NaCl, 1X Protease inhibitor cocktail). The homogenates were sonicated thrice by 5 seconds pulse with a 30-second gap in ice followed by the addition of an equal volume of protein sample loading buffer (2X) to the lysate and boiled for 3 minutes (at 100°C) in a dry bath. Protein concentration was estimated by Lowry method [17]. 20μg of larval lysate was subjected to 10%/15% SDS-PAGE and the expression of the respective proteins was determined by Western blot (Table S2). β-tubulin was used as the loading control. The blots were developed using SuperSignal West Pico Chemiluminescent Substrate (Pierce) and captured using ChemiDoc imaging system (Syngene). Densitometric analysis of protein bands was quantified using ImageJ software.

Starvation assay in WT, CG2135^-/-^ and glucose and 3-BDO fed CG2135^-/-^ larvae were done in 2% agar vials containing 1X PBS (Phosphate Buffer saline; pH-7.4) devoid of any fly food for 24 hrs.

### 2.5. *Drosophila* larval bodyweight and buoyancy screening

For overall gross estimation of the distribution of larval fat body of wildtype and CG2135^−/−^, age-matched larvae (40hrs A.E.L: after egg laying) of each genotype were collected and immobilized for 1hr at -20°C followed by capturing whole-body images in Apotome.2 microscope (Zeiss, Germany). For individual bodyweight experiments, age-matched larvae (120hrs A.E.L) of both WT and CG2135^−/−^ were measured using a fine balance. Buoyancy screening was performed in 10% sucrose solution as mentioned previously [18]. In brief, age-matched larvae of both WT and CG2135^−/−^ were transferred to 10% sucrose solution with a fine brush and allowed to attain equilibrium for 5 mins based on their buoyancy. The percentages of larvae floating and submerged for each were then measured individually both for WT and CG2135^−/−^ flies.

### 2.6. Larval fat body dissection and Nile Red staining of lipid droplets

Fat bodies of wandering 3^rd^ instar larva were dissected under a dissection microscope with Dumont (No.5) forceps in 1X Phosphate Buffer saline (PBS) (pH-7.4). The dissected fat tissues were fixed with 4% paraformaldehyde (PFA) for 30 mins at room temperature (RT) followed by washing with IX PBS and then incubated in dark for 30 mins in 1:2000 dilution of 0.5mg/ml Nile red stock solution (Sigma Aldrich). Subsequently, the tissues were rinsed with 1X PBS and then mounted using VectaShield mounting media containing DAPI (Vector Laboratories). All fluorescent images were taken in Olympus IX 81 Epiflourescense and Zeiss LSM 710 confocal microscopes and processed using the microscope’s software (ZEN). Quantification of individual areas of lipid droplets was done using ImageJ software.

### 2.7. Glucose and 3-BDO feeding to larvae

CG2135^−/−^ larvae (30 larvae each) were reared in 10g/l of glucose/200µM of 3BDO (3-Benzyl-5-((2-nitrophenoxy) methyl)-dihydrofuran-2(3H)-one) (Sigma Aldrich) supplemented in standard fly food. Post feeding, the larvae were starved in 2% agar vials containing 1X PBS devoid of any fly food for 24 hrs.

### 2.8. Imaging studies

Autophagosomal distribution of starved (24hrs) 3^rd^ instar larval fat tissues of WT and CG2135^-/-^ transgenic constructs were studied from orthogonal microscopic projections of z-stacked images captured in Apotome.2 microscope (Zeiss, Germany) at 40X oil immersion and processed using microscope’s software (ZEN). In brief, the fat body of wandering 3^rd^ instar starved transgenic larvae of both genotypes were dissected under a dissection microscope with Dumont (No.5) forceps in 1X Phosphate Buffer saline (PBS) (pH-7.4). The dissected fat tissues were then fixed with 4% paraformaldehyde (PFA) for 30 mins at room temperature (RT) followed by washing with IX PBS and mounted using VectaShield mounting media containing DAPI (Vector Laboratories). All fluorescent images were then captured in Apotome.2 microscope (Zeiss, Germany) at 40X oil immersion. Quantitative counts of colocalization points and the Pearson’s correlation coefficient (r) values were determined using the Colocalization module of ZEN software.

### 2.9. Statistical analysis

All statistical analysis was done from at least 3 independent experiments by Student’s t-test (2-tailed) using GraphPad Prism software (GraphPad Software Inc.) for comparison between two groups and One-way ANOVA for more than two groups and the values are expressed as Standard error means (SEM). P values indicate significance levels symbolized with an asterisk with *P≤0.05, **P≤0.01, ***P≤0.001 and ****P≤0.0001.

## 3. Results

### 3.1. CG2135^-/-^ fly exhibits lean phenotype due to paucity of adipose storage

Murine MPS VII was originally characterized by severe adipose storage deficiency along with significantly reduced body weight [4,5]. Lower body mass index (BMI) is also a common feature in many MPS VII patients [3]. However, the cause of this adipose deficiency or loss of body weight in MPS VII is not known. We attempted to understand the underlying mechanism of this intriguing phenomenon using a *Drosophila* model of MPS VII (CG2135^-/-^ fly) that we recently developed [14]. First, we analyzed the larval fat tissues, which are well dispersed and easily distinguishable at this stage. Gross microscopic observation revealed that as compared to their wild type counterparts, the CG2135^-/-^ larvae (40hrs AEL) were strikingly more transparent, which is indicative of adipose tissue atrophy (Fig. 1A). This observation closely resembles the lipid-deficient phenotype seen in dLipin mutant larvae or those overexpressing β-spectrin [19,20]. The CG2135^-/-^ larvae also showed ∼1.5-fold reduction in body weight that might be attributed to loss of adipose tissue mass (Fig. 1B). To further validate the loss of adipose tissue in the CG2135^-/-^ larvae, we employed a buoyancy-based assay. This assay is based on the principle that individuals with greater fat content float better in a solution of fixed density than individuals with lower fat content [18,21]. Strikingly, 84% of the wild type larvae were found to float on the surface of the sucrose solution as opposed to only 7% of the CG2135^-/-^ larvae (Fig. 1C, D). This data clearly indicated that the CG2135^-/-^ larvae had significantly reduced adipose storage compared to their wild type counterpart.

**Fig: 1.**
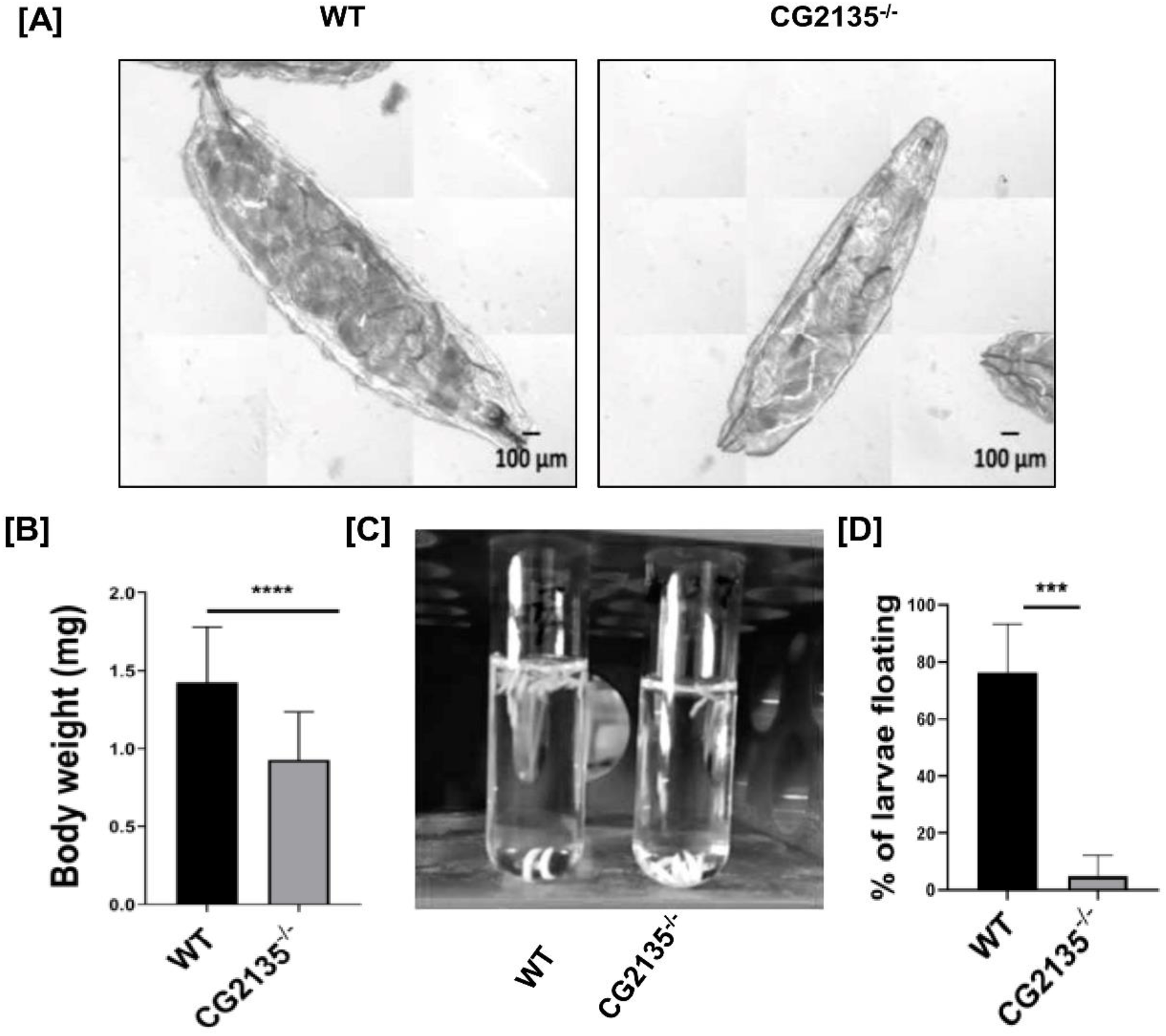
Reduced adipose storage and lean body mass in the CG2135^-/-^ larvae: (A) Age-matched whole body images displaying stored body fat of WT and CG2135^-/-^ larvae (40hrs A.E.L.) at 10X magnification (n=8, Scale bar=100μm). (B) Bar graph representing the individual body weight of WT and CG2135^-/-^ larvae at 120 hrs A.E.L. (WT n=84, CG2135^-/-^ n= 64). (C and D) Buoyancy assay in 10% sucrose solution and its representative bar diagram indicating 84% of WT and 7% of CG2135^-/-^ larvae floating on the surface (n=20 larvae each). Error bars represent the standard error mean (SEM). Asterisks represent a level of significance; ***P≤0.001 and ****P≤0.0001; Student’s t-test).

### 3.2. Fewer adipocytes, smaller lipid droplets and reduced adipogenesis in the CG2135^-/-^ fly

Since fat cell (adipocyte) number and lipid contents in them are the primary determinants of the overall fat storage, we next analyzed the adipocytes in the larval fat body by staining with DAPI (a nuclear stain) and Nile Red (a stain for the intracellular lipid droplets). Upon counting the DAPI stained nuclei, we found that the number of adipocytes in the CG2135^-/-^ larval fat body were ∼ 1.65 folds less than in the wild type larvae (Fig. 2A). As shown in Fig. 2B, the lipid droplets in the adipocytes were loosely packed (with several gaps between them, indicated by arrows) in the CG2135^-/-^ larval fat body as opposed to their compact organization in the wild type larvae. Interestingly, we also observed that while in the wild type larval fat body the lipid droplets were of uniform size, their sizes varied widely in the CG2135^-/-^ larvae. Upon further analysis, it was revealed that the average size of the lipid droplets in the CG2135^-/-^ larval fat body was considerably smaller (mean area: 23.8±11.5 μM^2^) as opposed to larger lipid droplets (of area 28.9±7.3 μM^2^) in the wild type larvae (Fig. 2C). Such small-sized lipid droplets were earlier found in the fat store deficient dLipin mutant fly [22]. We also checked for the expression of the adipogenic regulator, dSREBP/HLH106 and lipid droplet-associated proteins, Lipid storage droplet-1 and 2 (Lsd-1 and Lsd-2) in the adult flies [23–25]. It is worth mentioning that mutations in these genes were reported to cause a lean phenotype in flies [26,27]. It was observed that the transcript levels of dSREBP/HLH106 and Lsd-2 were reduced by 1.5 and 1.8-folds, respectively, in the CG2135^-/-^ flies as compared to their wild type counterparts (Fig S1:A,B). Taken together, our results suggest that reduced adipogenesis is responsible for adipose storage deficiency in the CG2135^-/-^ fly. Dissecting the underlying reason behind this phenotype was our next objective.

**Fig: 2.**
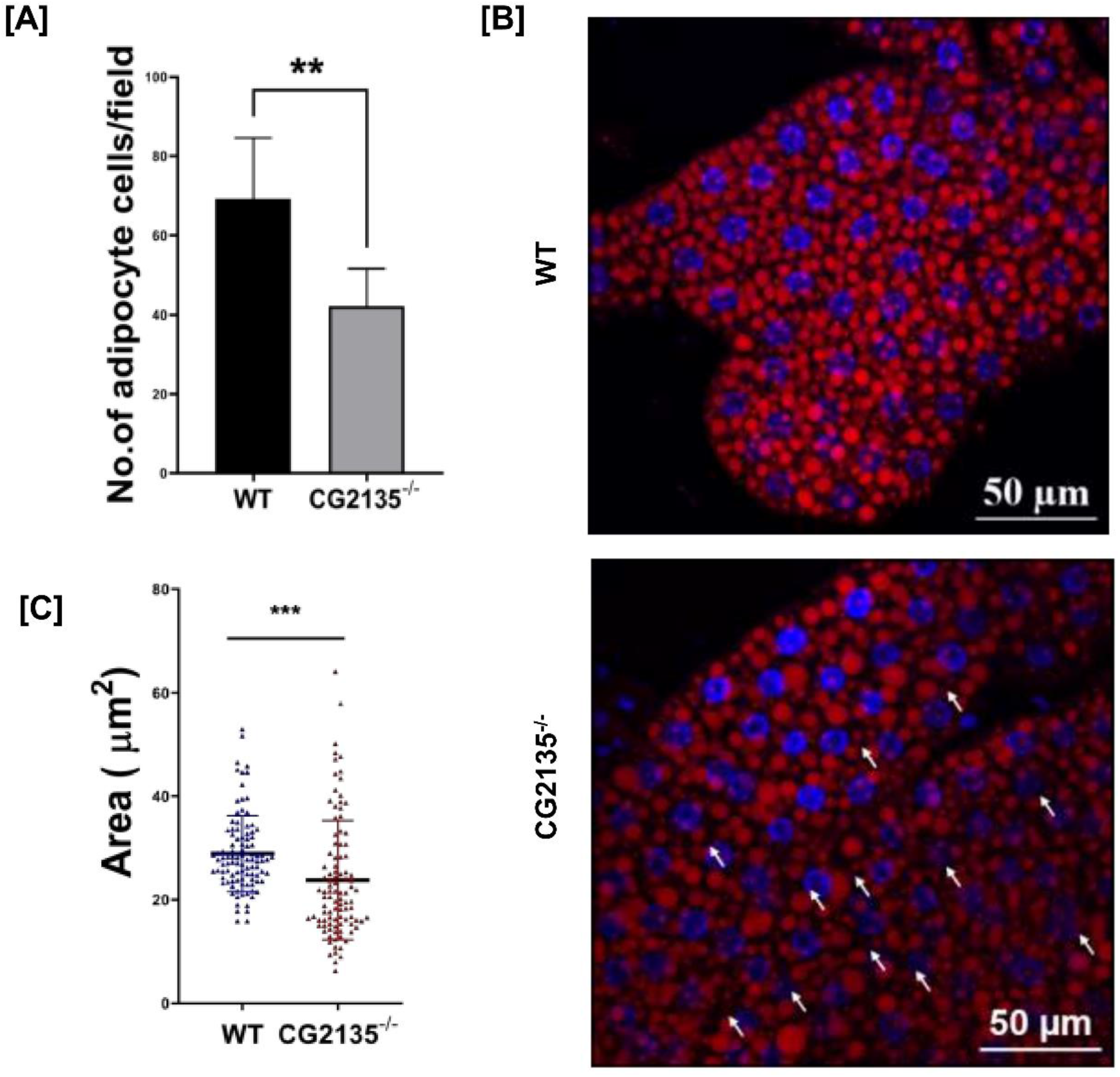
Perturbed adipogenesis in the CG2135^-/-^ flies. (A) The numbers of adipocyte nuclei post staining of lipid globules with DAPI were quantitated both for the WT and CG2135^-/-^ larvae (WT: n=7; CG2135^-/-^: n=6) (B) Nile Red stained dissected fat tissues of wandering 3^rd^ instar WT and CG2135^-/-^ larvae at 40X magnification (Scale bar=50µm). The arrows in the CG2135^-/-^ larval panel represent deformed morphology. The gaps within the lipid droplets also indicate less compact distribution in CG2135^-/-^ larvae on being compared to WT larvae (Blue: DAPI stained nuclei; Red: Nile Red stained lipid globules). (C) The individual area of lipid globules of both genotypes was measured using ImageJ software (WT n=100, CG2135^-/-^ n= 100 lipid droplets) showing a 1.21-fold significant reduction for CG2135^-/-^ larvae as compared to its wildtype counterpart. Error bars represent standard error mean (SEM). Asterisks represent a level of significance (**P≤0.01;***P≤0.001; Student’s t-test).

### 3.3. Abnormal autophagy due to autophagosome-lysosome fusion defect in the CG2135^-/-^ fly

A direct relationship between impaired autophagy and reduced fat storage has been reported earlier through elegant genetic studies [12,13]. It was shown that adipose-specific deletion of the important autophagy-related gene (*atg7*) resulted in drastically reduced adipogenesis in mice [13]. These reports, together with the fact that autophagy impairment has emerged as a common feature in many lysosomal storage disorders [28,29], prompted us to check the status of autophagy in the CG2135^-/-^ fly. For this, we starved the larvae for 24 hours to induce autophagy and then analyzed the status of Atg8a (fly homolog of the autophagy marker, LC3) by western blot. Data presented in Fig3A, B shows that the Atg8a-II level in the CG2135^-/-^ larvae were significantly elevated (∼3.8 folds) as compared to the wild type flies, suggesting an increased abundance of autophagosomes in the mutant flies. To assess whether this increased accumulation of autophagosomes in the CG2135^-/-^ larvae is due to stimulated autophagy pathway or impaired autophagosome-lysosome fusion, we over-expressed GFP-mCherry-Atg8a transgene by *Actin*-GAL4 driver in the wildtype (CG2135^+/+^) and the CG2135^-/-^ genetic background independently (Fig S2:A, B). [15]. In these flies, the double fluorophore tagged Atg8 protein is supposed to emit yellow fluorescence (green and red merged) in non-acidic autophagosomal compartments and only red fluorescence in the autolysosomes (autophagosome fused with lysosomes) where the green signal from GFP gets quenched off due to acidic pH [15]. Genomic DNA-PCR was performed with CG2135 amplifying primers to validate the genetic background of the transgenic flies (Fig S2:C). Overall, as compared to the wild type, we observed a significant increase in yellow fluorescence in the fat body of the CG2135^-/-^ larvae under starved conditions (Fig. 3C). This increased yellow fluorescence in the CG2135^-/-^ larval fat body resulted from a significant amount of green (GFP) and red (mCherry) colocalization as indicated by Pearson’s correlation coefficient values (Fig. 3D). This data suggests that there is an increased abundance of starvation-induced autophagosomes in the CG2135^-/-^ larval fat body, which is due to autophagosome-lysosome fusion defect. A similar impairment in the autophagic flux was earlier seen in the chondrocytes of the β-glucuronidase knockout mouse [30]. Our data indicate that defective autophagy in the fat body might be responsible for the adipose storage deficiency in the CG2135^-/-^ fly.

**Fig: 3.**
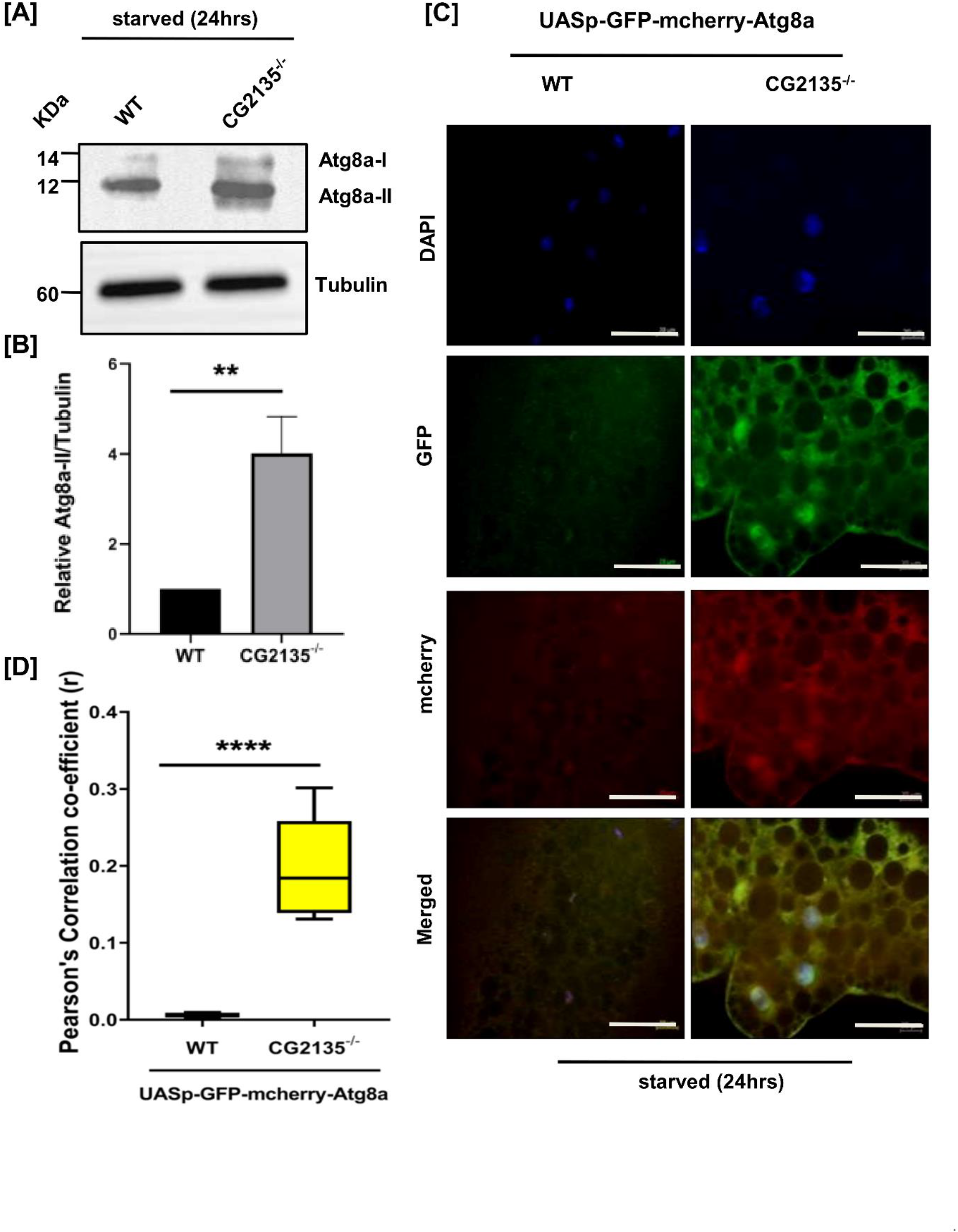
Aberrant autophagosome-lysosome fusion in fat tissues of the CG2135^-/-^ larvae: (A) Protein level of Atg8a [Atg8a-I: 14KDa: Atg8a-II: 12KDa) in starved (24hrs) larval lysate (20μg) of WT and CG2135^-/-^ was checked by western blotting using an antibody against *Drosophila* LC3 homolog (Atg8a). Expression of β-tubulin (60KDa), detected by anti-β-tubulin antibody was considered as the loading control (B) Graph comparing relative Atg8a-II band intensity (12KDa) of WT and CG2135^-/-^ larvae. (C) Orthogonal microscopic fluorescent images of z-stacked fat tissues depicting the distribution of autophagosomes (yellow vacuoles) in starved 3^rd^ instar larval fat tissues of both UASp-GFP-mcherry-Atg8a-(WT and CG2135^-/-^) transgenic construct driven by Actin-GAL4 at 40X magnification (Scale bar=20μm). (DAPI stained nuclei are shown in blue, Atg8a-GFP expression in green and Atg8a-mcherry expression in red). (D) Quantitative counts of the colocalization points measured from the Pearson’s correlation coefficient (r) values of UASp-GFP-mcherry-Atg8a-WT and UASp-GFP-mcherry-Atg8a-CG2135^-/-^ transgenic larvae post 24hrs starvation are represented in the Whisker’s box plot. The range of r are indicated as follows: UASp-GFP-mcherry-Atg8a-WT: r = 0.001333-0.00928; UASp-GFP-mcherry-Atg8a-CG2135^-/-^: r = 0.13103-0.30161. (UASp-GFP-mcherry-Atg8a-WT: n=5 larval fat tissues, UASp-GFP-mcherry-Atg8a-CG2135^-/-^: n=8 larval fat tissues). Error bars represent the standard error mean (SEM). Asterisks represent a level of significance (**P≤0.01; ****P≤0.0001; Student’s t-test).

### 3.4. Autophagy defect and adipose deficiency in the CG2135^-/-^ larvae is corrected by treatment with the mTOR stimulator 3BDO

We next explored whether defective autophagy and the adipose deficiency phenotype in the CG2135^-/-^ fly can be reversed by pharmacological intervention. Since autophagy-lysosomal pathway and mTOR signaling are reciprocally regulated, we focused our attention to this intriguing network as a possible target [31,32]. It is well-established that mTOR acts as a negative regulator of autophagy and rapamycin-mediated inhibition of mTOR has been reported to induce autophagy in various cell types (Fig 4A) [33]. Conversely, pharmacological impairment of lysosome function that mimics lysosomal storage resulted in feedback inhibition of mTOR activity [34,35]. This was thought to be a cellular response to tackle lysosome malfunction by upregulating the autophagic capacity of the cell. We were thus prompted to check for mTOR activity in our flies by measuring the phosphorylation status of its downstream target S6K [36]. Western blot with p70S6K (Thr^398^) antibody revealed more than 2 folds reduction in S6K phosphorylation in the CG2135^-/-^ larvae as compared to their wild type counterparts (Fig 4B, C). Our data thus confirmed that mTOR activity is indeed downregulated in the CG2135^-/-^ flies as has been observed in some other models of lysosomal storage diseases [37,38]. This data provided the rationale for treating the flies with mTOR stimulator and check if the autophagy defect and adipose deficiency in the CG2135^-/-^ fly can be corrected. Nutrients such as glucose and amino acids are generic activators of mTOR [39]. Recently, Ge *et. al*. identified 3BDO [3-Benzyl-5-((2-nitrophenoxy) methyl)-dihydrofuran-2(3H)-one] as a novel and specific mTOR activator [40]. Hence, we attempted to restore mTOR activity in the CG2135^-/-^ larvae by rearing them in glucose (10g/l) or 3BDO (200µM) containing fly food. Indeed, treatment with these mTOR stimulators restored the phosphorylation status of S6K in the CG2135^-/-^ flies as revealed by our western blot data (Fig.4D, E). Importantly, treatment with 3BDO or glucose significantly reduced the Atg8a-II titer in the CG2135^-/-^ larvae to the wild type level (Fig. 4F, G, S4. A, B). This data indicates that pharmacological stimulation of mTOR activity in the CG2135^-/-^ larvae were successful in correcting abnormal upregulation of autophagy in those flies. Whether 3BDO can replenish the adipose deficiency in the CG2135^-/-^ larvae was tested next. Interestingly, CG2135^-/-^ larvae fed with 3BDO showed significant restoration of their body weight to the level almost comparable to their wild type counterparts (Fig. 5A). When subjected to buoyancy assay, the 3BDO-fed CG2135^-/-^ larvae also showed a remarkable increase in the percentage of floaters (>80%) compared to their counterparts reared on normal fly food (only 7% floaters) (Fig. 5B, C). We also analyzed the effect of 3BDO treatment on the lipid droplet size in CG2135^-/-^ larval fat body. It was observed that the average area of lipid droplets in the 3BDO-fed CG2135^-/-^ larval fat body was 34.21±13.93μm^2^ as opposed to only 22.07±11.43μm^2^ and 28.63±6.96μm^2^ for the untreated CG2135^-/-^ or wild type larvae, respectively (Fig. 5D, E). A similar restoration of the adipose storage in the CG2135^-/-^ larvae was also observed upon glucose feeding (Fig. S3:A-E). Collectively, these data are testimony of the fact that pharmacological stimulation of mTOR activity indeed corrected the adipose storage deficiency phenotype of the CG2135^-/-^ larvae.

**Fig: 4.**
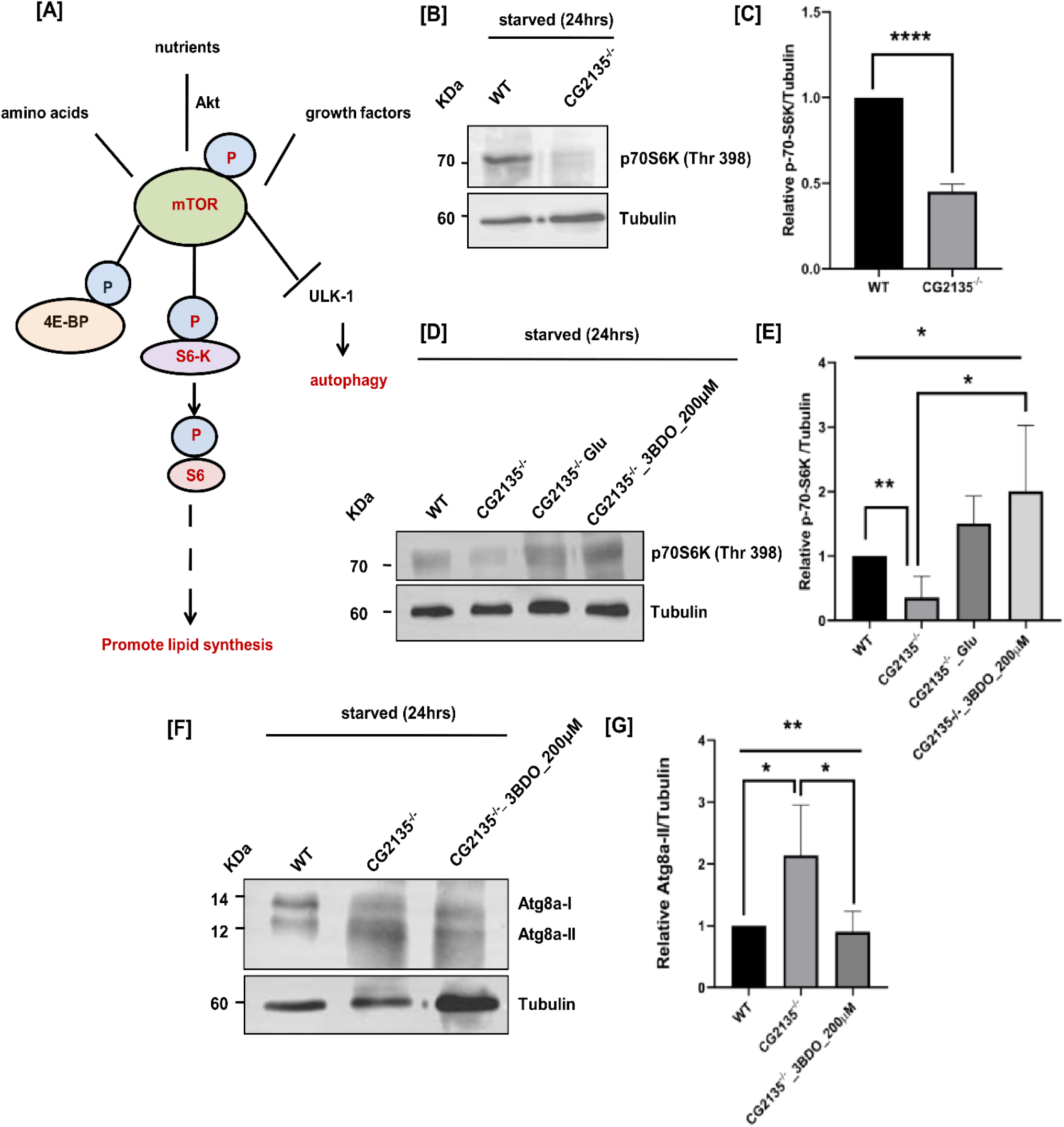
Pharmacological stimulation of mTORC1 activity in CG2135^-/-^ larvae corrects autophagy defect: (A) Schematic representation showing positive stimulators of mTORC1 such as nutrients (glucose), amino acids, and growth factors. One of the downstream targets of mTORC1 is p-S6K which on activation promotes lipid synthesis and impairs autophagy by inhibition of the ULK-1 complex (Unc-51 activating kinase-1) (B) Immunoblotting of WT and CG2135^-/-^ starved larval lysate (24hrs) with *Drosophila* specific p70S6K (Thr398) antibody (70KDa). 20µg of the starved lysates were loaded in each well and β-tubulin (60KDa) was used as the loading control. (C) Bar graph comparing relative p70S6K (Thr398) (70KDa) band intensity of WT and CG2135^-/-^ larvae. (D) Western blotting of starved larval lyaste with *Drosophila* p70S6K (Thr398) antibody (70KDa) post-supplementation of glucose (10g/l) / 3BDO (200µM) to CG2135^-/-^ larvae. 20µg of the respective protein lysates were loaded in each well and β-tubulin (60KDa) was used as the loading control (E) Representative bar diagram depicting the relative p70S6K (Thr398) (70KDa) band intensity of WT, CG2135^-/-^, CG2135^-/-^_Glucose and CG2135^-/-^_3BDO larvae post starvation (24hrs). (F) Protein level of Atg8a [Atg8a-I: 14KDa: Atg8a-II: 12KDa) in starved (24hrs) larval lysate (20μg) of WT, CG2135^-/-^ and CG2135^-/-^ 3BDO (200 µM) was checked by western blotting using *Drosophila* Atg8a antibody. Expression of β-tubulin (60KDa), detected by anti-β-tubulin antibody was considered as the loading control (G) Graph comparing relative Atg8a-II band intensity (12KDa) of WT, CG2135^-/-^ and 3BDO supplemented CG2135^-/-^ (200 µM) larvae. Error bars represent standard error mean (SEM). Asterisks represent a level of significance (\**P*≤0.05, **P≤0.01, ***P≤0.001, ****P≤0.0001; Student’s t-test and One-way ANOVA).

**Fig: 5.**
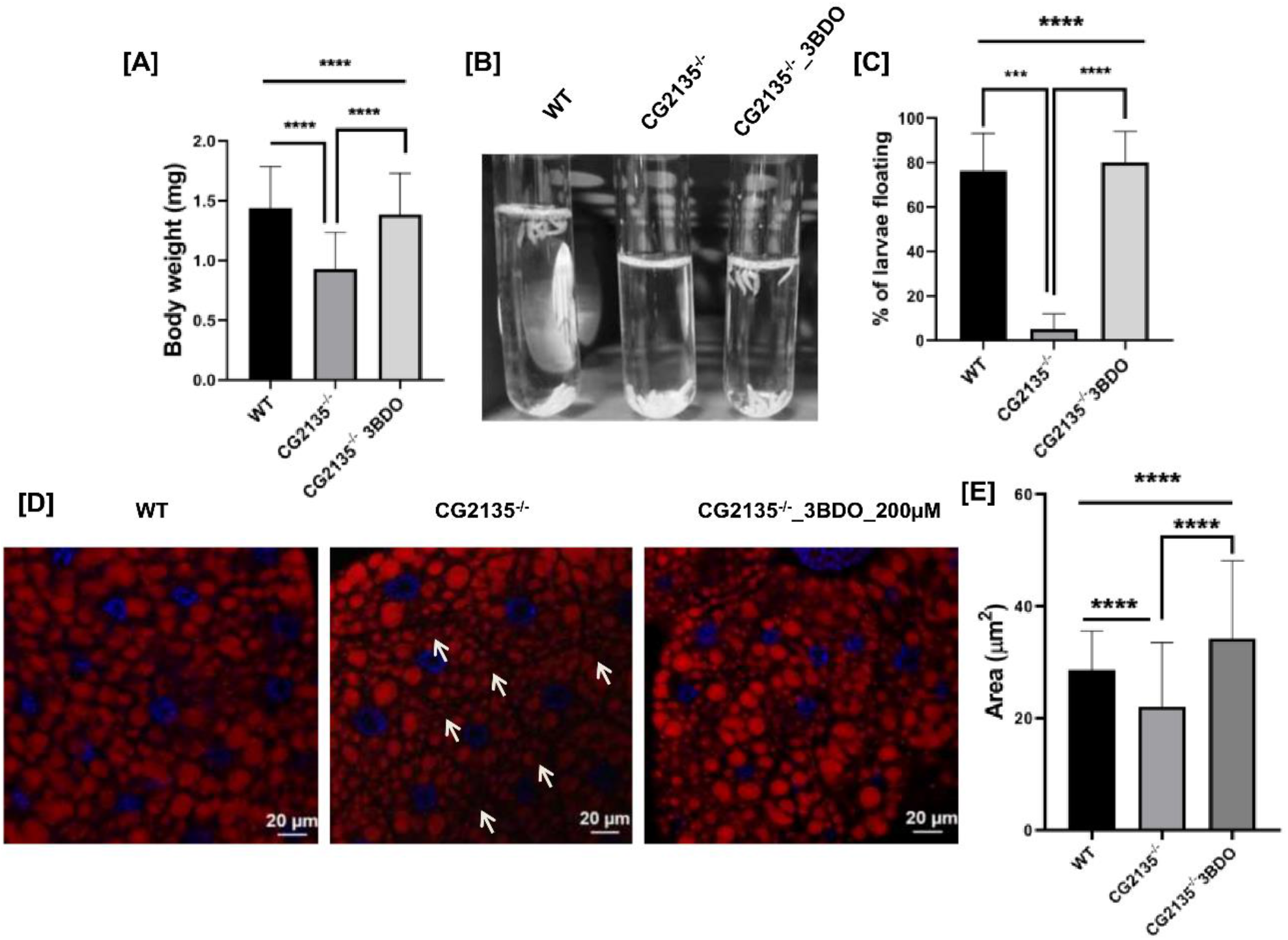
3BDO treatment restores adipose storage in the CG2135^-/-^ larvae: (A) Bar diagram representing the individual body weight of 3^rd^ instar (120hrs A.E.L.) WT, CG2135^-/-^ and 3-BDO supplemented CG2135^-/-^ larvae (200 µM) (WT: n=88, CG2135^-/-^: n=63, CG2135^-/-^ 3BDO: n=39). (B) Recovery of the stored body fat in CG2135^-/-^ larvae was determined from the buoyancy assay post-administration with 3-BDO (200 µM) (C) Representative bar graph indicating the percentages of larvae floating for WT, CG2135^-/-^ and 3-BDO fed 3^rd^ instar CG2135^-/-^ larvae in 10% sucrose solution (D) Nile Red stained dissected fat tissues of wandering 3^rd^ instar WT, CG2135^-/-^ and CG2135^-/-^ 3BDO (200 µM) larvae depicting the arrangement of the lipid globules at 40X magnification (Scale bar=20µm). Loosely packed distribution and empty spaces in CG2135 ^-/-^ larval lipid droplets are demarcated by arrows (Blue: DAPI stained nuclei; Red: Nile Red stained lipid globules) (E) The individual area of lipid droplets are quantitated and represented in the bar graph for the WT, CG2135 ^-/-^ and CG2135^-/-^ 3BDO (200 µM) larval fat body (WT: n=100, CG2135^-/-^: n=100, CG2135^-/-^ 3BDO: n=74 lipid droplets). Error bars represent standard error mean (SEM). Asterisks represent a level of significance (\**P*≤0.05, **P≤0.01, ***P≤0.001 and ****P≤0.0001; Student’s t-test and One-way ANOVA).

## 4. Discussion

Creation of the first *Drosophila* model of MPS VII by knocking out the fly β-GUS gene (CG2135) was a breakthrough in MPS VII research. This fly model recapitulated cardinal features of the disease including neuromuscular degeneration and progressive loss of locomotor function [14]. Our continued exploration of this novel fly model now revealed that similar to the β-GUS deficient mice, the CG2135^-/-^ flies also exhibited a lean phenotype with significantly reduced adipose storage. This allowed us to investigate the mechanism of reduced adiposity in MPS VII. We report here that adipose deficiency in the CG2135^-/-^ flies is due to reduced adipogenesis which might be a consequence of aberrant autophagy and stunted mTOR activity observed in these mutant flies.

Our striking finding that autophagy defect and adipose deficiency in the CG2135^-/-^ flies could be corrected by treating them with mTOR stimulators pointed towards a hitherto unknown mechanistic link between mTOR activity, autophagy pathway and adipogenesis in MPS VII. Significantly reduced bodyweight of the CG2135^-/-^ larvae was attributed to adipose deficiency, which in turn resulted from a fewer number of adipocytes and lesser lipid contents in them. Signs of impaired adipogenesis were also apparent in the adult CG2135^-/-^ flies. These data although explained the lean phenotype of the MPVII patients or the β-GUS deficient MPS VII mouse model, raised further questions about the underlying mechanism behind the loss of adiposity. It was earlier observed that functional autophagy plays a critical role in the adipogenic process and genetic disruption of autophagy resulted in a gross reduction in white adipose tissue mass in mice [12,13]. So, it was important for us to examine the status of autophagy in the CG2135^-/-^ flies, which eventually revealed an increased abundance of autophagosomes owing to autophagosome-lysosome fusion defect in their larval fat body. This observation corroborated with several other prior studies reporting dysfunctional autophagy in patient-derived fibroblast cells or animal models of MPSIIIA, multiple sulfatase deficiency, neuronal ceroid lipofuscinoses (NCLs), Gaucher disease and Pompe disease [29,41–43]. Impairment of autophagy and accumulation of autophagosomes and autophagic substrates was also observed in MPS VII primary chondrocyte cells [30]. Thus, it seemed quite plausible that impaired autophagy is the key reason behind reduced adipogenesis in the CG2135^-/-^ flies.

Our finding that mTOR activity is downregulated in the CG2135^-/-^ larvae is reminiscent of the perturbed mTOR signaling observed in the muscle of the Pompe disease mouse model [37]. This reduction in mTOR activity is most likely a feedback mechanism to upregulate the autophagy-mediated turnover capacity of the cell for compensating the loss of lysosome function [34,35]. However, in the CG2135^-/-^ larvae this compensatory mechanism might be detrimental because it would lead to abnormal accumulation of autophagosomes in the cytosol owing to the autophagosome-lysosome fusion defect seen in these flies. This notion is corroborated by our finding that the abundance of autophagosomes was found to be indeed high in the CG2135^-/-^ larval fat body. This may induce cytotoxicity in the adipocytes as has been reported earlier in other cell types [44]. Since mTOR is a negative regulator of autophagy, we hypothesized that stimulation of mTOR activity could prevent abnormal accumulation of autophagosomes and may restore adipose storage in the CG2135^-/-^ flies [31]. It is worth mentioning that in Pompe mouse model dampened mTOR activity was genetically restored by knocking down the TSC2 gene, a negative regulator of mTOR. This resulted in the clearance of abnormally accumulated autophagosomes and prevented muscle degeneration [37]. Instead of taking a genetic approach, we decided to tweak mTOR activity in the CG2135^-/-^ flies using its pharmacological activators, 3BDO and glucose. Complete absence of autophagic buildup and restored adipose storage in the 3BDO or glucose-fed CG2135^-/-^ larva provided testimony for our hypothesis. Results with 3BDO are particularly encouraging since it specifically targets the mTOR activation pathway and its therapeutic efficacy, bioavailability, as well as safe dosage, have already been established in a murine model of Alzheimer’s disease [45]. 3BDO treatment may improve the metabolic health of the MPS VII patients since adipose deficiency is known to trigger several metabolic syndromes [46,47]. Whether treatment of the CG2135^-/-^ flies with 3BDO can provide any other therapeutic benefits like life span extension or attenuation of locomotor defect is an exciting question that remains to be investigated. The present study opens up the possibility of exploring mTOR-directed therapy in MPS VII.

## Conclusion

Collectively, our results elucidate the mechanism of adipose deficiency in MPS VII, which was poorly understood to date. Abnormal autophagy due to defect in autophagosome-lysosome fusion, coupled with downregulation of mTOR activity constitutes a vicious cycle that ultimately resulted in reduced adipogenesis in the CG2135^-/-^ flies. This mechanistic insight provided a crucial lead for therapeutic intervention to correct the lean phenotype of the CG2135^-/-^ flies with mTOR activators, glucose and 3BDO. Knowledge generated from this study may have far-reaching implications in understanding the molecular basis of adipose storage deficiency in other lysosomal storage disorders.

## Abbreviations

A.E.L.: after egg laying
GAGs: glycosaminoglycans
WT: wildtype
KO: knockout
Atg8a: Autophagy related 8a
LC3: Microtubule-associated protein 1A/1B light chain 3B
mTORC1: Mammalian target of rapamycin complex 1
MPS VII: Mucopolysaccharidosis Type VII
LSD: Lysosomal storage disorder
β-GUS: β-glucuronidase
S6K: Ribosomal protein S6 kinase beta
3BDO: 3-Benzyl-5-((2-nitrophenoxy) methyl)-dihydrofuran-2(3H)-one.

## Acknowledgements

The authors sincerely thank Mr. Ritabrata Ghosh and Mr. Sujoy Bose for their expert technical assistance. The authors are thankful to Drs. Jayasri Das Sarma and Sankar Maiti of IISER Kolkata for their help with various reagents and constructive suggestions. This research was supported by the Department of Biotechnology (DBT) and Ministry of Education, Government of India grants BT/PR6423/GBD/27/437/2012 and STARS/APR2019/BS/779/FS, respectively. Fly maintenance was supported by the Department of Biotechnology (DBT) and Ministry of Education, Government of India grants BT/PR5775/BRB/10/1466/2015 and STARS/APR2019/BS/591 respectively to MP. IB was supported by IISER Kolkata Ph.D. fellowship and SB was supported by University Grants Commission fellowship.

## Supplementary information

**Table S1:**
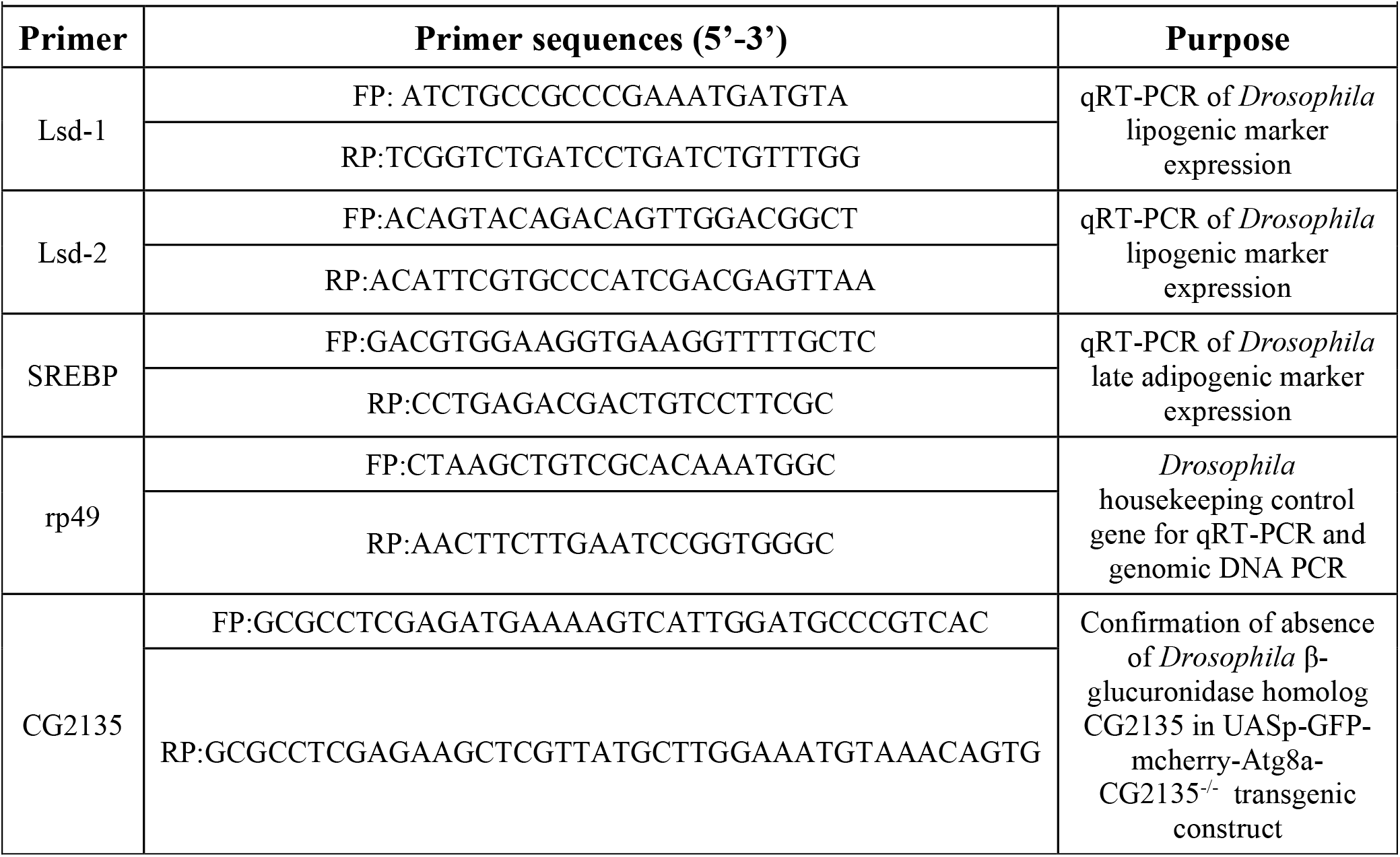
Oligonucleotides used in this work.

**Table S2:** Antibodies used in this work.

| Table S2: Antibodies used in this work |  |  |
| --- | --- | --- |
| Primary Antibodies | Dilution used | Source |
| Mouse polyclonal anti β-Tubulin antibody (T9026) | 1:5000 | Sigma Aldrich |
| Rabbit polyclonal anti –ATG8 antibody | 1:2000 | Merck |
| Rabbit polyclonal anti-p70S6K (Thr 398) antibody | 1:1000 | PhosphoSolutions |
| Secondary Antibodies | Dilution used | Source |
| Anti-Rabbit IgG (whole molecule)-Peroxidase | 1:4000 | Sigma Aldrich |
| Anti-Mouse IgG (whole molecule)-Peroxidase | 1:4000 | Sigma Aldrich |

**FigS1.**
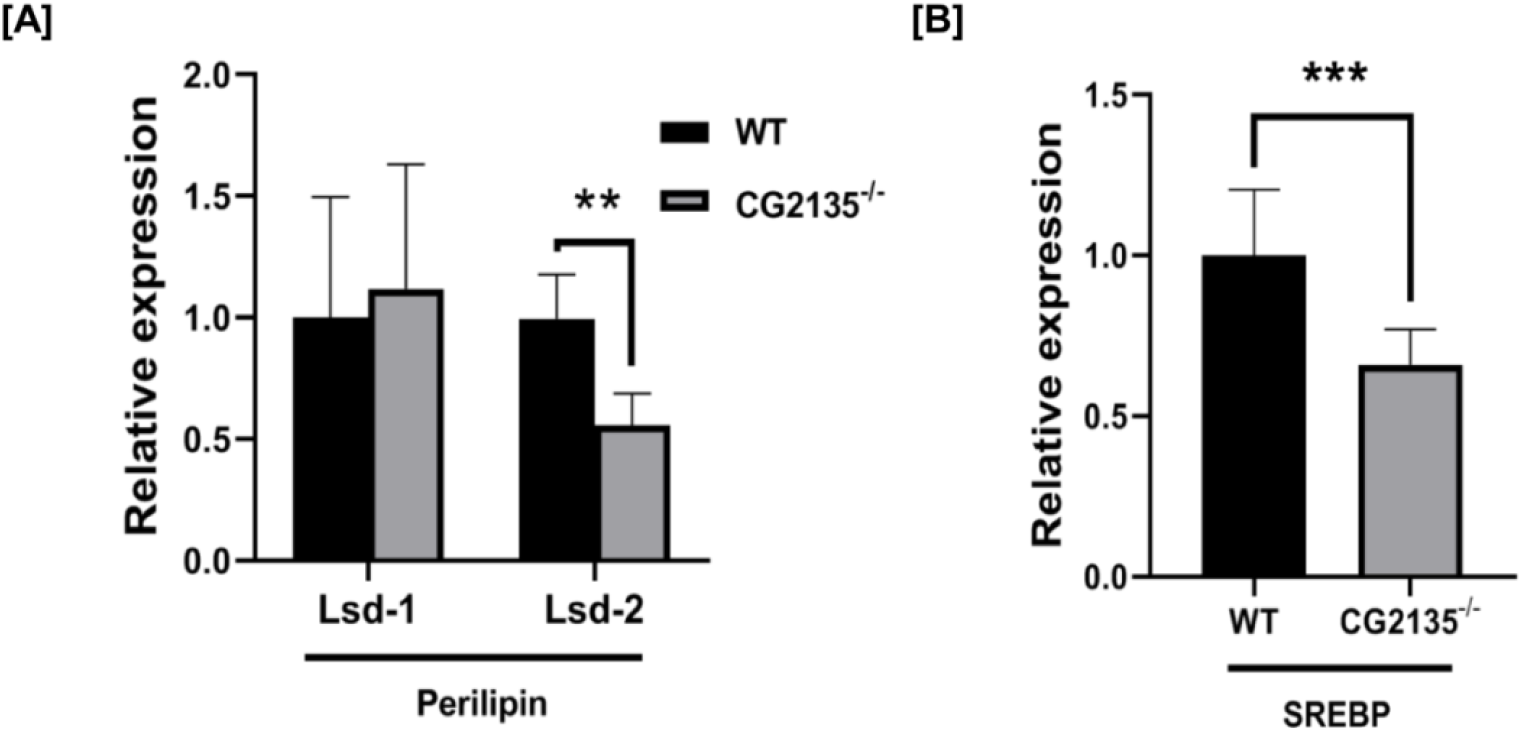
Aberrant adipogenesis in CG2135^-/-^ flies: qRT-PCR showing relative expression pattern of adipogenic markers in WT and CG2135^-/-^ flies (8-day old adult flies). CG2135^-/-^ flies showed (A) 1.8-fold significant downregulation in the relative expression of Lsd-2 (*Drosophila* Perilipin homolog) and (B) 1.5-fold significant reduction in expression of SREBP as compared to WT flies. rp49 was used as the reference control gene. Error bars represent standard error mean (SEM). Asterisks represent a level of significance (\**P*≤0.05, **P≤0.01 and ***P≤0.001; Student’s t-test).

**FigS2.**
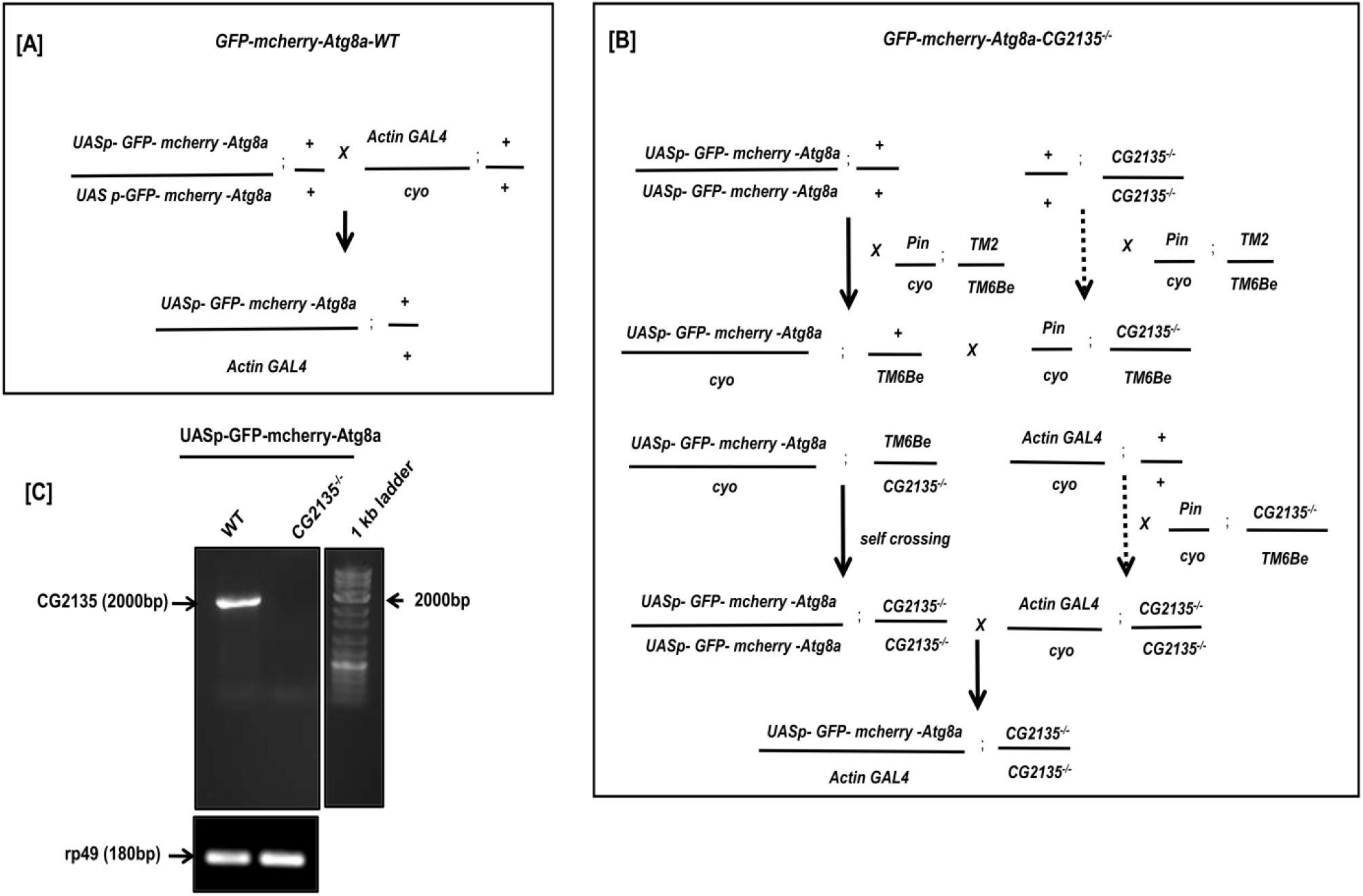
Generation and affirmation of UASp-GFP-mcherry-Atg8a transgenic construct flies w.r.t. WT and CG2135^-/-^ background: Schematic representation depicting the cross-setup for generation of: (A) UASp-GFP-mCherry-Atg8a-WT transgenic construct: This construct was generated by crossing y[1]w[1118];P{w[+mC]=UASp-GFP-mCherry-Atg8a}2 flies with W; Actin GAL4/cyo driver and (B) UASp-GFP-mCherry-Atg8a-CG2135^-/-^ construct flies was obtained by bringing the y[1]w[1118];P{w[+mC]=UASp-GFP-mCherry-Atg8a}2 fly line with respect to the CG2135^-/-^ background. Independent crosses were set for both the fly lines with balancer flies, Pin/Cyo;TM2/TM6Be. After selection of the desired constructs in each round of crossing, the UASp-GFP-mcherry-Atg8a line was brought in the background of CG2135^-/-^ under the control of Actin5C-GAL4 driver. (C) Validation by genomic DNA-PCR of UASp-GFP-mCherry-Atg8a transgenic construct (WT and CG2135^−/−^) background using respective forward and reverse primers for CG2135 gene (2000bp). rp49 (180bp) was used as the housekeeping control gene.

**FigS3.**
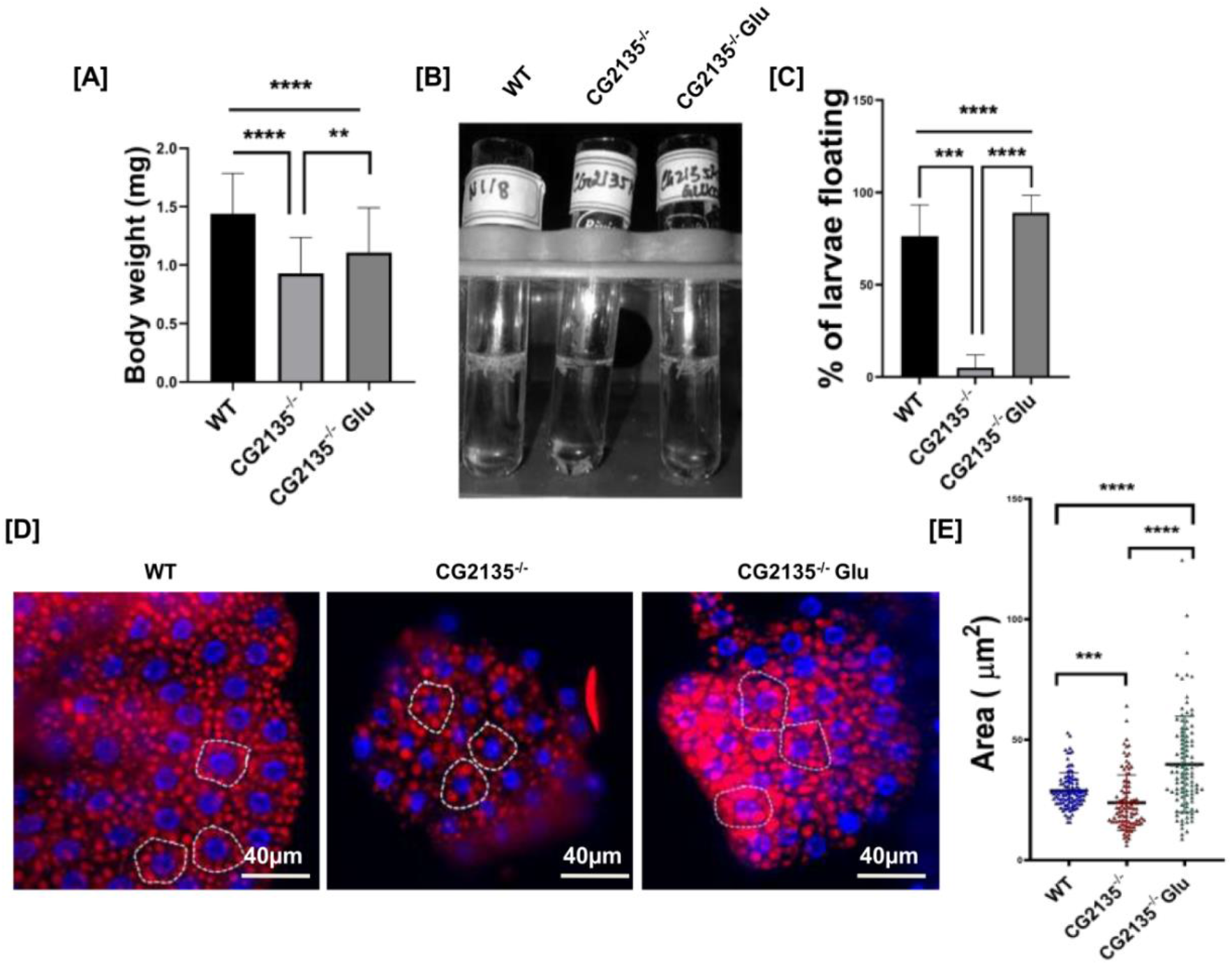
mTORC1 stimulation with glucose rescues adipose deficiency in CG2135^−/−^ larvae: (A) Bar graph representing the individual body weight of 3^rd^ instar (120 hrs A.E.L.) WT, CG2135^-/-^ and glucose supplemented CG2135^-/-^ larvae (WT: n=88, CG2135^-/-^ : n=63, CG2135^-/-^ Glu: n=66). (B,C) Bouyancy screening of WT, CG2135^-/-^ and glucose fed 3^rd^ instar CG2135^-/-^ larvae showing percentages of larvae floating in 10% sucrose solution (D) Nile Red stained dissected fat tissues of wandering 3^rd^ instar WT, CG2135^-/-^ and CG2135^-/-^ Glu larvae depicting the arrangement of the lipid globules at 40X magnification (Scale bar=40µm). Distribution of the clone of larval lipid droplets in WT, CG2135 ^-/-^ and CG2135^-/-^ Glu are demarcated by white dotted grids (Blue: DAPI stained nuclei; Red: Nile Red stained lipid globules) (E) The individual area of lipid globules are quantitated and represented in the bar graph for the WT, CG2135 ^-/-^ and CG2135^-/-^ Glu larval fat tissues (WT: n=100, CG2135^-/-^: n=100, CG2135^-/-^ Glu: n=100 lipid droplets). Error bars represent standard error mean (SEM). Asterisks represent a level of significance (\**P*≤0.05, **P≤0.01, ***P≤0.001 and ****P≤0.0001; Student’s t-test and One-way ANOVA).

**FigS4.**
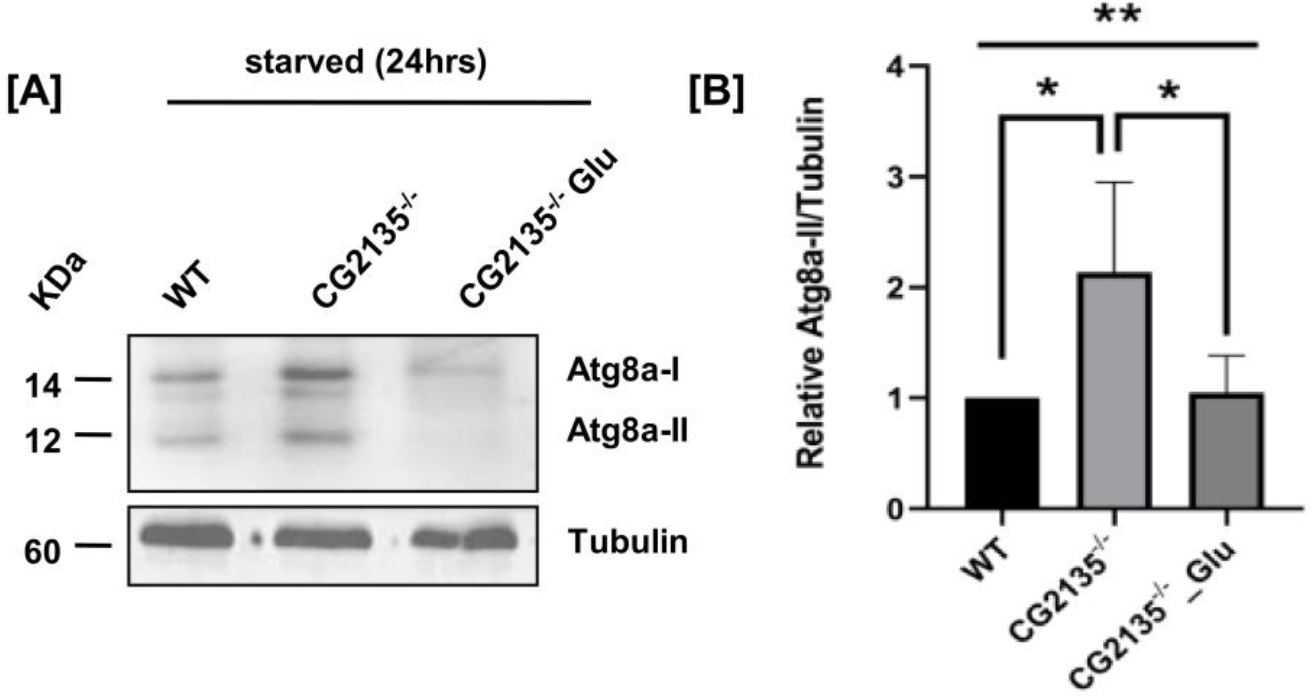
Amelioration of defective autophagic flux in glucose supplemented CG2135^-/-^ larvae: (A) Immunoblotting of starved WT, CG2135^-/-^ and glucose fed CG2135^-/-^ larval lysate with *Drosophila* LC3 (Atg8a) antibody. 20μg of the respective starved larval lysates were loaded in each well and β-tubulin (60KDa) was used as the loading control. (B) Graph comparing relative Atg8a-II band intensity (12KDa) of WT, CG2135^-/-^ and Glucose supplemented CG2135^-/-^ larvae. Error bars represent standard error mean (SEM). Asterisks represent a level of significance (\**P*≤0.05 and **P≤0.01; Student’s t-test and One-way ANOVA).

## References

[1] S. WS, Q. BA, M. WH, R. DL, Beta glucuronidase deficiency: report of clinical, radiologic, and biochemical features of a new mucopolysaccharidosis, The Journal of Pediatrics. 82 (1973) 249–257. https://doi.org/10.1016/S0022-3476(73)80162-3.

[2] P. G, A. G, B. A, Lysosomal storage diseases: from pathophysiology to therapy, Annual Review of Medicine. 66 (2015) 471–486. https://doi.org/10.1146/ANNUREV-MED-122313-085916.

[3] M. AM, L.-H. N, S. RD, G. BH, S. M, G. R, P. M, G.-M. A, Ç. M, B. D, S. MS, W. R, G. R, M. A, P. G, K. AK, E. F, T. A, A. L, B. M, F. RE, B. K, F. J, W. KK, M. GA, C. L, H. M, S. WS, Clinical course of sly syndrome (mucopolysaccharidosis type VII), Journal of Medical Genetics. 53 (2016) 403–418. https://doi.org/10.1136/JMEDGENET-2015-103322.

[4] B. EH, D. MT, B. WG, G. RE, V. CA, G. B, L. KA, M. LM, W. CJ, Murine mucopolysaccharidosis type VII. Characterization of a mouse with beta-glucuronidase deficiency, The Journal of Clinical Investigation. 83 (1989) 1258–1266. https://doi.org/10.1172/JCI114010.

[5] J.C. Woloszynek, T. Coleman, C.F. Semenkovich, M.S. Sands, Lysosomal Dysfunction Results in Altered Energy Balance, Journal of Biological Chemistry. 282 (2007) 35765–35771. https://doi.org/10.1074/JBC.M705124200.

[6] A.L. Ghaben, P.E. Scherer, Adipogenesis and metabolic health, Nature Reviews Molecular Cell Biology 2018 20:4. 20 (2019) 242–258. https://doi.org/10.1038/s41580-018-0093-z.

[7] D. Moseti, A. Regassa, W.-K. Kim, Molecular Regulation of Adipogenesis and Potential Anti-Adipogenic Bioactive Molecules, International Journal of Molecular Sciences 2016, Vol. 17, Page 124. 17 (2016) 124. https://doi.org/10.3390/IJMS17010124.

[8] S.R. Farmer, Transcriptional control of adipocyte formation, Cell Metabolism. 4 (2006) 263–273. https://doi.org/10.1016/J.CMET.2006.07.001.

[9] W. JC, K. A, O. KK, R. M, S. MS, Metabolic adaptations to interrupted glycosaminoglycan recycling, The Journal of Biological Chemistry. 284 (2009) 29684–29691. https://doi.org/10.1074/JBC.M109.020818.

[10] M. Clemente-Postigo, A. Tinahones, R. el Bekay, M.M. Malagón, F.J. Tinahones, The Role of Autophagy in White Adipose Tissue Function: Implications for Metabolic Health, Metabolites. 10 (2020). https://doi.org/10.3390/METABO10050179.

[11] S. R, H. Y, R. Y, N. M, R. A, L. G, Autophagy in adipose tissue and the beta cell: implications for obesity and diabetes, Diabetologia. 57 (2014) 1505–1516. https://doi.org/10.1007/S00125-014-3255-3.

[12] R. Singh, Y. Xiang, Y. Wang, K. Baikati, A.M. Cuervo, Y.K. Luu, Y. Tang, J.E. Pessin, G.J. Schwartz, M.J. Czaja, Autophagy regulates adipose mass and differentiation in mice, The Journal of Clinical Investigation. 119 (2009) 3329–3339. https://doi.org/10.1172/JCI39228.

[13] Y. Zhang, S. Goldman, R. Baerga, Y. Zhao, M. Komatsu, S. Jin, Adipose-specific deletion of autophagy-related gene 7 (atg7) in mice reveals a role in adipogenesis, Proceedings of the National Academy of Sciences. 106 (2009) 19860–19865. https://doi.org/10.1073/PNAS.0906048106.

[14] S. Bar, M. Prasad, R. Datta, Neuromuscular degeneration and locomotor deficit in a Drosophila model of mucopolysaccharidosis VII is attenuated by treatment with resveratrol, Disease Models & Mechanisms. 11 (2018). https://doi.org/10.1242/DMM.036954.

[15] I.P. Nezis, B. v. Shravage, A.P. Sagona, T. Lamark, G. Bjørkøy, T. Johansen, T.E. Rusten, A. Brech, E.H. Baehrecke, H. Stenmark, Autophagic degradation of dBruce controls DNA fragmentation in nurse cells during late Drosophila melanogaster oogenesis, Journal of Cell Biology. 190 (2010) 523–531. https://doi.org/10.1083/JCB.201002035.

[16] K.J. Livak, T.D. Schmittgen, Analysis of Relative Gene Expression Data Using Real-Time Quantitative PCR and the 2−ΔΔCT Method, Methods. 25 (2001) 402–408. https://doi.org/10.1006/METH.2001.1262.

[17] L. OH, R. NJ, F. AL, R. RJ, Protein measurement with the Folin phenol reagent, The Journal of Biological Chemistry. 193 (1951) 265–275. https://pubmed.ncbi.nlm.nih.gov/14907713/ (accessed August 31, 2021).

[18] T. Reis, M.R. van Gilst, I.K. Hariharan, A Buoyancy-Based Screen of Drosophila Larvae for Fat-Storage Mutants Reveals a Role for Sir2 in Coupling Fat Storage to Nutrient Availability, PLOS Genetics. 6 (2010) e1001206. https://doi.org/10.1371/JOURNAL.PGEN.1001206.

[19] U. R, L. Y, P. J, S. S, L. M, Lipin is a central regulator of adipose tissue development and function in Drosophila melanogaster, Molecular and Cellular Biology. 31 (2011) 1646–1656. https://doi.org/10.1128/MCB.01335-10.

[20] B. Diaconeasa, G.H. Mazock, A.P. Mahowald, R.R. Dubreuil, Genetic Studies of Spectrin in the Larval Fat Body of Drosophila melanogaster: Evidence for a Novel Lipid Uptake Apparatus, Genetics. 195 (2013) 871–881. https://doi.org/10.1534/GENETICS.113.155192.

[21] K.E. Hazegh, T. Reis, A Buoyancy-based Method of Determining Fat Levels in Drosophila, Journal of Visualized Experiments : JoVE. 2016 (2016) 54744. https://doi.org/10.3791/54744.

[22] R. Ugrankar, Y. Liu, J. Provaznik, S. Schmitt, M. Lehmann, Lipin Is a Central Regulator of Adipose Tissue Development and Function in Drosophila melanogaster, Molecular and Cellular Biology. 31 (2011) 1646–1656. https://doi.org/10.1128/MCB.01335-10.

[23] T. U, E. S, H. D, HLH106, a Drosophila transcription factor with similarity to the vertebrate sterol responsive element binding protein, Proceedings of the National Academy of Sciences of the United States of America. 93 (1996) 1195–1199. https://doi.org/10.1073/PNAS.93.3.1195.

[24] J. Bi, Y. Xiang, H. Chen, Z. Liu, S. Grönke, R.P. Kühnlein, X. Huang, Opposite and redundant roles of the two Drosophila perilipins in lipid mobilization, Journal of Cell Science. 125 (2012) 3568–3577. https://doi.org/10.1242/JCS.101329.

[25] P.E. Bickel, J.T. Tansey, M.A. Welte, PAT proteins, an ancient family of lipid droplet proteins that regulate cellular lipid stores, Biochimica et Biophysica Acta. 1791 (2009) 419. https://doi.org/10.1016/J.BBALIP.2009.04.002.

[26] A.S. Kunte, K.A. Matthews, R.B. Rawson, Fatty acid auxotrophy in Drosophila larvae lacking SREBP, Cell Metabolism. 3 (2006) 439–448. https://doi.org/10.1016/J.CMET.2006.04.011.

[27] T. L, R. C, R. P, E. A, V. NF, Drosophila Perilipin/ADRP homologue Lsd2 regulates lipid metabolism, Mechanisms of Development. 120 (2003) 1071–1081. https://doi.org/10.1016/S0925-4773(03)00158-8.

[28] L. AP, P. R, R. N, S. S, W. SU, B. A, Autophagy in lysosomal storage disorders, Autophagy. 8 (2012) 719–730. https://doi.org/10.4161/AUTO.19469.

[29] S. C, F. A, J. L, S. C, V. C, M. D, de P. R, T. C, R. DC, B. A, A block of autophagy in lysosomal storage disorders, Human Molecular Genetics. 17 (2008) 119–129. https://doi.org/10.1093/HMG/DDM289.

[30] B. R, C. L, D.L. C, F. A, S. AC, M. J, D.G. E, N. E, A. I, L. C, S. M, L. B, B. A, S. C, mTORC1 hyperactivation arrests bone growth in lysosomal storage disorders by suppressing autophagy, The Journal of Clinical Investigation. 127 (2017) 3717–3729. https://doi.org/10.1172/JCI94130.

[31] D.-T. S, P.-P. ME, F. FJ, C. JL, The role of TOR in autophagy regulation from yeast to plants and mammals, Autophagy. 4 (2008) 851–865. https://doi.org/10.4161/AUTO.6555.

[32] K. YC, G. KL, mTOR: a pharmacologic target for autophagy regulation, The Journal of Clinical Investigation. 125 (2015) 25–32. https://doi.org/10.1172/JCI73939.

[33] J. Kim, M. Kundu, B. Viollet, K.-L. Guan, AMPK and mTOR regulate autophagy through direct phosphorylation of Ulk1, Nature Cell Biology 2011 13:2. 13 (2011) 132–141. https://doi.org/10.1038/ncb2152.

[34] L. M, K. B, Z. H, K. JH, C. X, C. D, V. L, L. PQ, V. A, Y. XM, Suppression of lysosome function induces autophagy via a feedback down-regulation of MTOR complex 1 (MTORC1) activity, The Journal of Biological Chemistry. 288 (2013) 35769–35780. https://doi.org/10.1074/JBC.M113.511212.

[35] A.O. Fedele, C.G. Proud, Chloroquine and bafilomycin A mimic lysosomal storage disorders and impair mTORC1 signalling, Bioscience Reports. 40 (2020) 20200905. https://doi.org/10.1042/BSR20200905.

[36] M.K. Holz, J. Blenis, Identification of S6 Kinase 1 as a Novel Mammalian Target of Rapamycin (mTOR)-phosphorylating Kinase*, Journal of Biological Chemistry. 280 (2005) 26089–26093. https://doi.org/10.1074/jbc.M504045200.

[37] J.-A. Lim, L. Li, O.S. Shirihai, K.M. Trudeau, R. Puertollano, N. Raben, Modulation of mTOR signaling as a strategy for the treatment of Pompe disease, EMBO Molecular Medicine. 9 (2017) 353–370. https://doi.org/10.15252/EMMM.201606547.

[38] O.B. Davis, H.R. Shin, C.-Y. Lim, R.M. Perera, M.P. Ordonez, R.Z. Correspondence, NPC1-mTORC1 Signaling Couples Cholesterol Sensing to Organelle Homeostasis and Is a Targetable Pathway in Niemann-Pick Type C Graphical Abstract Highlights d Proteomic profiling of NPC lysosomes reveals both proteolytic and structural defects d Loss of cholesterol transport activity by NPC1 causes aberrant mTORC1 signaling d mTORC1 inhibition restores lysosomal and mitochondrial function in NPC cells, Developmental Cell. 56 (2021) 260-276.e7. https://doi.org/10.1016/j.devcel.2020.11.016.

[39] J.L. Jewell, K.-L. Guan, Nutrient Signaling to mTOR and Cell Growth, Trends in Biochemical Sciences. 38 (2013) 233. https://doi.org/10.1016/J.TIBS.2013.01.004.

[40] D. Ge, L. Han, S. Huang, N. Peng, P. Wang, Z. Jiang, J. Zhao, L. Su, S. Zhang, Y. Zhang, H. Kung, B. Zhao, J. Miao, Identification of a novel MTOR activator and discovery of a competing endogenous RNA regulating autophagy in vascular endothelial cells, Autophagy. 10 (2014) 957. https://doi.org/10.4161/AUTO.28363.

[41] K. M, S. M, W. S, Y. K, T. I, K. E, G. T, P. C, von F. K, M. N, S. P, U. Y, Participation of autophagy in storage of lysosomes in neurons from mouse models of neuronal ceroid-lipofuscinoses (Batten disease), The American Journal of Pathology. 167 (2005) 1713–1728. https://doi.org/10.1016/S0002-9440(10)61253-9.

[42] K. KJ, G. S, C.-Q. JI, W. NS, L. L, S. E, G. M, M. K, H. J, B. I, P. L, A Drosophila Model of Neuronopathic Gaucher Disease Demonstrates Lysosomal-Autophagic Defects and Altered mTOR Signalling and Is Functionally Rescued by Rapamycin, The Journal of Neuroscience: The Official Journal of the Society for Neuroscience. 36 (2016) 11654–11670. https://doi.org/10.1523/JNEUROSCI.4527-15.2016.

[43] Y. Cao, J.A. Espinola, E. Fossale, A.C. Massey, A.M. Cuervo, M.E. MacDonald, S.L. Cotman, Autophagy Is Disrupted in a Knock-in Mouse Model of Juvenile Neuronal Ceroid Lipofuscinosis, Journal of Biological Chemistry. 281 (2006) 20483–20493. https://doi.org/10.1074/JBC.M602180200.

[44] B. RW, R. SL, W. TL, H. CO, L. S, Accumulation of autophagosomes confers cytotoxicity, The Journal of Biological Chemistry. 292 (2017) 13599–13614. https://doi.org/10.1074/JBC.M117.782276.

[45] W. L, Y. H, X. Z, Y. S, Y. H, Z. C, W. P, X. S, M. J, Z. B, B. J, A butyrolactone derivative 3BDO alleviates memory deficits and reduces amyloid-β deposition in an AβPP/PS1 transgenic mouse model, Journal of Alzheimer’s Disease: JAD. 30 (2012) 531–543. https://doi.org/10.3233/JAD-2012-111985.

[46] L. Scheja, J. Heeren, The endocrine function of adipose tissues in health and cardiometabolic disease, Nature Reviews Endocrinology 2019 15:9. 15 (2019) 507–524. https://doi.org/10.1038/s41574-019-0230-6.

[47] S.M. Grundy, Adipose tissue and metabolic syndrome: too much, too little or neither, Eur J Clin Invest. 45 (2015) 1209–1217. https://doi.org/10.1111/eci.12519.

